# Adaptive parallel tempering for BEAST 2

**DOI:** 10.1101/603514

**Authors:** Nicola F. Müller, Remco R. Bouckaert

## Abstract

With ever more complex models used to study evolutionary patterns, approaches that facilitate efficient inference under such models are needed. Parallel tempering has long been used to speed up phylogenetic analyses and to make use of multi-core CPUs. Parallel tempering essentially runs multiple MCMC chains in parallel. All chains are heated except for one cold chain that explores the posterior probability space like a regular MCMC chain. This heating allows chains to make bigger jumps in phylogenetic state space. The heated chains can then be used to propose new states for other chains, including the cold chain. One of the practical challenges using this approach, is to find optimal temperatures of the heated chains to efficiently explore state spaces. We here provide an adaptive parallel tempering scheme to Bayesian phylogenetics, where the temperature difference between heated chains is automatically tuned to achieve a target acceptance probability of states being exchanged between individual chains. We first show the validity of this approach by comparing inferences of adaptive parallel tempering to MCMC on several datasets. We then explore where parallel tempering provides benefits over MCMC. We implemented this adaptive parallel tempering approach as an open source package licensed under GPL 3.0 to the Bayesian phylogenetics software BEAST2, available from https://github.com/nicfel/CoupledMCMC.

## Introduction

Phylogenetic methods are being used to study increasingly complex processes. Analyses using such methods, however, also require an increasingly large amount of computational resources. One way still be able to perform these analyses is by making use of multiple CPU’s, which requires calculations to be parallelisable. Tree likelihood calculations (Suchard and Rambaut, 2009) often assume independent evolutionary processes on different branch and nucleotide sites and can be easily parallelised (Suchard and Rambaut, 2009). This can, however, be complex or even impossible for many other parts of such analyses, most notably tree prior calculations, which are used to infer demographic processes from phylogenetic trees. A lot of recent development in the filed of phylogentics has been focused on developing such tree priors that allow us to infer complex population dynamics from genetic sequence data. As a result, analyses using standard Bayesian tools such as Markov chain Monte Carlo (MCMC), can be very time consuming. This, in turn, limits the datasets that can be studied and the complexity of models that can be used to do so.

Alternatively, parallel tempering can be used to speed up analyses in Bayesian phylogenetics (Altekar et al., 2004). This approach is based on running multiple MCMC chains, each at a different “temperature”, which effectively flattens the posterior probability space. This allows heated chains to move faster through the posterior probability space, and increases the chance to travel between local optimas. After some amount of iterations, two chains are randomly exchanged in what is essentially an MCMC move. In such a move, the parameters of the two chains are exchanged, but each chain keeps its temperatures. While the heated chains do not explore the true posterior probabilities, the one cold chain does. In contrast to MCMC, however, parallel tempering requires additional parameters to setup an analysis. Defining the temperatures of each chain in particular, can be problematic and may require some amount of testing. Choosing sub-optimal temperatures of chains can lead to inefficient exploration of the posterior probability space, essentially wasting the additional computational resources used.

The problem of finding good temperatures is related to the issue of finding good variances of proposal distributions in MCMC. One way to deal with that is to automatically adapt variances in proposal distributions to achieve optimal acceptance probabilities of moves during an MCMC (Haario et al., 2001). This can be applied to adaptively tune the temperatures of heated chains in the parallel tempering framework (Miasojedow et al., 2013). We here employ this adaptive mechanism to tuning the temperature difference between chains in the parallel tempering algorithm. To do so, we update the temperature difference between chains, whose temperatures are geometrically distributed (Kofke, 2002), during a parallel tempering run. The amount by which the temperature is updated is increasingly being reduced during each run, which eventually leads the temperatures of chains to be approximately constant (Haario et al., 2001). While not being Markovian, this leads the algorithm to be ergodic.

We implemented this adaptive parallel tempering algorithm in BEAST 2 (Bouckaert et al., 2014), where a lot of novel Bayesian phylogenetic model development takes place (Bouckaert et al., 2019). This implementation makes use of multiple CPU cores, allowing virtually any analyses in BEAST 2 to be performed on multi-core machines increasing the size of datasets that can be analyzed and the complexity of models that can be used to do so. By default, the implementation adapts the temperature difference between heated chains to achieve an acceptance probability of any two chains being exchanged of 0.234 (Roberts et al., 1997, 2001; Kone and Kofke, 2005; Atchadé et al., 2011).

We first show the correctness of the adaptive parallel tempering approach by comparing summary statistics of multi type tree distributions sampled under the structured coalescent (Vaughan et al., 2014) to the summary statistics received when using regular MCMC. Additionally, we show that distributions of posterior probability estimates are constant over the course of analyses using adaptive parallel tempering, when inferring past population dynamics of Hepatitis C in Egypt (Ray et al., 2000; Pybus et al., 2003).

Next, we show how automatically tuning the temperature, leads to an acceptance probability that converges to the target probability from different initial temperatures on two different datasets.

We then compare MCMC to adaptive parallel tempering using different levels of heating on two different datasets. First, we apply it to the Hepatitis C dataset, where we do not expect regular MCMC to be stuck in local optimas. Then, we apply it to a dataset which has been described to be easily stuck in local optimas (Lakner et al., 2008; Hohna and Drummond, 2011).

## Methods and Material

### Background

Parallel tempering makes use of running *n* different chains *i* = 1,…,*n* at different temperatures (Geyer, 1991; Gilks and Roberts, 1996; Altekar et al., 2004). Each of the different chains works similar to a regular MCMC chain. In regular MCMC, a parameter space is explored as follows: Given that the MCMC is currently at state *x*, we propose a new state *x*’ from a proposal distribution *g*(*x*’|*x*) given the current state. At this new state, we calculate the likelihood *P*(*D*|*x*’) of the data *D* given the state and the prior probability of the new state *P*(*x*’) and compare it the to old state. The acceptance probability of accepting this new state is then calculated as follows:

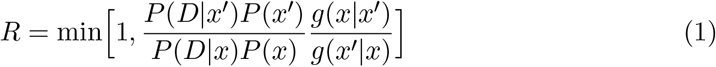

If *R* is greater than a randomly drawn value between [0,1], the new state *x*’ is accepted as the current state, otherwise it is rejected and we remain in the same state. If we keep proposing new states *x*’ and accept these using equation (1), we eventually explore parameter space with the frequency at which values of a parameter are visited being its marginal probability (Geyer, 1991).

One of the issues of using this approach is that acceptance probabilities can be quite low, which makes it hard to move between different states in parameter space. Alternatively, an MCMC chain can be heated by using a temperature scaler 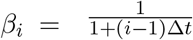, with *i* being the number of the chain (Altekar et al., 2004). Heating of an MCMC chain changes its acceptance probability *R_heated_* to:

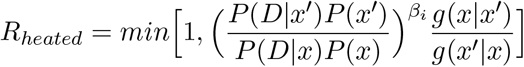

For a heated chain however, the frequency at which a value of a parameter is visited does not correspond to its marginal probability any more. However, heated chains can be used as a proposal to update the cold chain by performing what is essentially an MCMC move. This move proposes to swap the current states of two random chains i and *j* with the temperature *β_i_* and *β_j_* such that *β_i_* < *β_j_*. Exchanging the states of chains *i* and *j* is accepted with an acceptance probability *R_ij_* of:

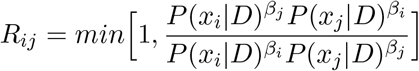

As for a regular MCMC move, swapping the states of the two chains is accepted when a randomly drawn uniformly distribution value in [0,1] is smaller than *R_ij_*.

### Locally aware adaptive tuning of the temperature of heated chains

Choosing an optimal temperature of the different heated chains can be a tedious task, requiring running an analysis, updating temperatures of the analysis and re-running everything. Instead, the temperatures of chains can be tuned automatically during the run itself to achieve a targeted average acceptance probability. As stated above, we consider the temperatures of *n* different chains to be geometrically distributed and the tune the temperature difference Δ*t* during the analysis.

When updating the temperature based on the global acceptance probability, we compute *p_current_* based on all proposed exchanges of states from the start of a run to the current state. We then iteratively tune the temperature to achieve the target average acceptance probability *p_target_* over the course of an analysis as follows. At each proposed exchange of states between states, we denote the probability of an exchange being accepted as *p_current_*. Given *p_current_* and *p_target_,* we update the difference in temperature between chains Δ*t* as follows:

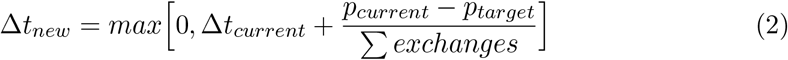

With Σ *exchanges* denoting the total number of proposed exchanges, which increases throughout the BEAST run. This means that updating the temperature as in equation (2), leads the tuning of the temperature to become smaller and smaller and eventually become 0.

Tuning Δ*t* is only performed after an initial burn-in period of (by default) 100 proposed exchanges. By default, the target acceptance probability is set to 0.234, which for many MCMC proposals can be shown to be an optimal trade-off between as many accepted moves as possible and as large of a move as possible (Kone and Kofke, 2005; Atchadé et al., 2011). Datasets where unfavourable intermediate states are of particular issue may, however, require higher temperatures and therefore lower acceptance probabilities to overcome these intermediate states.

Changing the temperature of a heated chain changes the equilibrium distribution of that chain. There can be a significant time lag between changing the temperature of a chain and that chain moving to its new equilibrium state. If the temperature is updated too fast, heated chains did not necessarily reach this new equilibrium, which in turn can lead to over-adaptation. This is particularly problematic at the beginning of an analysis where *exchanges* is relatively small and where large changes in the temperature could occur. In order to reduce the risk of that, we maximize the difference between Δ*t_current_* and Δ*t_new_* to be 0.001.

Another issue can arise when the global acceptance probability strongly differs from the current acceptance probability. In order to avoid that, we made the adaptation procedure aware of the local acceptance probability. To do so, we additionally compute a local acceptance probability *p_local_* of the last 100 proposed exchanges. We only update the temperature if the global and the local acceptance are on the same side of the target acceptance probability, i.e. if *p_locai_* > *p_target_* & *P_globai_* > *p_target_* or *p_local_* < *p_target_* & *p_global_* < *p_target_*.

### Implementation

In this implementation of the parallel tempering algorithm, we run *n* different MCMC chains, with each chain *i* ∈ [1,…,*n*] running at a temperature 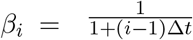. The temperatures of the different chains are therefore geometrically distributed, which has been shown to be a good spacing of temperatures between individual chains (Kofke, 2002).

Upon initialisation, we first sample at random at which iteration the states of two chains with which number are proposed to be exchanged. We then initialise each chain to be run in its own Java thread using multiple CPU cores, if available. Each chain is then run until it reaches the time when an exchange of states with another chain will be proposed. This means than every chain runs independently of each other until an iteration at which it actually participates in a proposed exchange, minimising the crosstalk between threads Altekar et al. (2004). If the exchange of states between different chains is accepted, we exchange the temperature of the two chains instead of the states themselves. The states can be quite large and exchanging them across different chains is potentially quite time consuming. Instead of exchanging the states themselves, we exchange the operator specifications and logger. Exchanging the operator specifications is done such that the individual tuning parameters of operators of a chain can be optimized to run at specific temperatures. The loggers are exchanged such that each heated chain logs its states to the log file that corresponds to its temperature and not the number of the chain.

The temperature is adapted at any potential exchange of states between chains, after an initial phase of 100 potential exchanges without any adaption. The temperature is updated simultaneously on all chains, not just the ones participating in the exchange of states, independent of which iterations they are in.

Adaptive parallel tempering is implemented, such that runs that were prematurely stopped or didn’t reach sufficient convergence yet can be resumed. Usually, a graphical user interface called BEAUti is used to set up BEAST 2 analyses. Setting up analyses with parallel tempering works differently depending on whether a BEAUTi template is needed to set up an analysis as required for some packages. If no such template is needed, an analysis can be set up to run with parallel tempering directly in BEAUTi and we provide a tutorial on how to do this on https://taming-the-beast.org/tutorials/CoupledMCMC-Tutorial/ (Barido-Sottani et al., 2017). Alternatively, we provide an interface that converts BEAST2 xmls setup to run with MCMC into such that run with adaptive parallel tempering.

## Data Availability and Software

The BEAST 2 package coupledMCMC can be downloaded by using the package manager in BEAUti. The source code for the software package can be found here: https://github.com/nicfel/CoupledMCMC. The XML files used for the analysis performed here can be found in https://github.com/nicfel/CoupledMCMC-Material. All plots were done using ggplot2 (Wickham, 2016) in R (Team et al., 2013).

## Validation

Similar to the validation of MCMC operators, we can sample under the prior to validate the implementation of the parallel tempering approach. To do so, we sampled typed trees with 5 taxa and two different states under the structured coalescent using Multi-TypeTree (Vaughan et al., 2014). We did this sampling once using MCMC and once using parallel tempering. If the implementation of the parallel tempering algorithm explores the same parameter space as MCMC, marginal parameter distributions sampled using both approaches should be equal. In figure S1, we compare the distribution of different summary statistics of typed trees between MCMC and parallel tempering.

## Results

### Ergodicity of the adaptive parallel tempering algorithm

First, we test if the distribution of posterior probability values using adaptive parallel tempering algorithm are consistent over time, i.e. ergodic. To do so, we ran 100 skyline (Drummond et al., 2005) analyses of Hepatitis C in Egypt (Ray et al., 2000), with 3 different target acceptance probabilities, 0.234 (Kone and Kofke, 2005; Atchadé et al., 2011), 0.468 (= 2 * 0.234) and 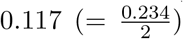. The temperature difference between chains Δ*t* is being adapted during the analyses, particularly during the initial phase (see figure 1A).

We then computed the distribution of posterior probability estimates of the 100 different runs using the posterior probability estimates at different iterations. The distribution of posterior probability estimates stays constant over the different iterations (see figure 1A), despite the temperature difference between chains being adapted. This is true for all 3 different target acceptance probabilities.

### Automatic tuning of the temperature of heated chains

We next tested how well the adaptive tuning of the temperature of heated chains over the course of an analysis works starting from different initial values. To do so, we ran to different datasets, the Hepatitis C dataset (Ray et al., 2000) as well an influenza A/H3N2 analysis using MASCOT as analyzed previously (Müller et al., 2018). We ran each dataset with 4 different initial temperatures (0.0001, 0.001, 0.01 and 0.1), each targeting 3 different acceptance probabilities, 0.234, 0.468 and 0.117. Additionally, we used two different frequencies to propose swaps between chains, once proposing swaps every 100 iterations and once every 1000. Since the temperature is adapted at every possible swap, this means that the runs with swaps every 100 iterations adapt Δ*t* 10 times more frequently than the ones proposing swaps every 1000 iterations. We kept the temperature scaler constant for the first 100 potential swaps of states between chains.

As shown in figure S2, for any of the here considered initial values of the temperature scaler, the target acceptance probability is reached quite early in the run and very well approximated at the end of the run using the Hepatitis C example. The same applies to the analysis of the influenza A/H3N2 dataset (see figure S4).

After an initial phase where the adaption of the temperature difference can overshoot the optimal value, Δ*t* is adapted such that it approximates the target value better and better during the run (see figures 2 and S3 for the MASCOT analysis).

**Figure 1:**
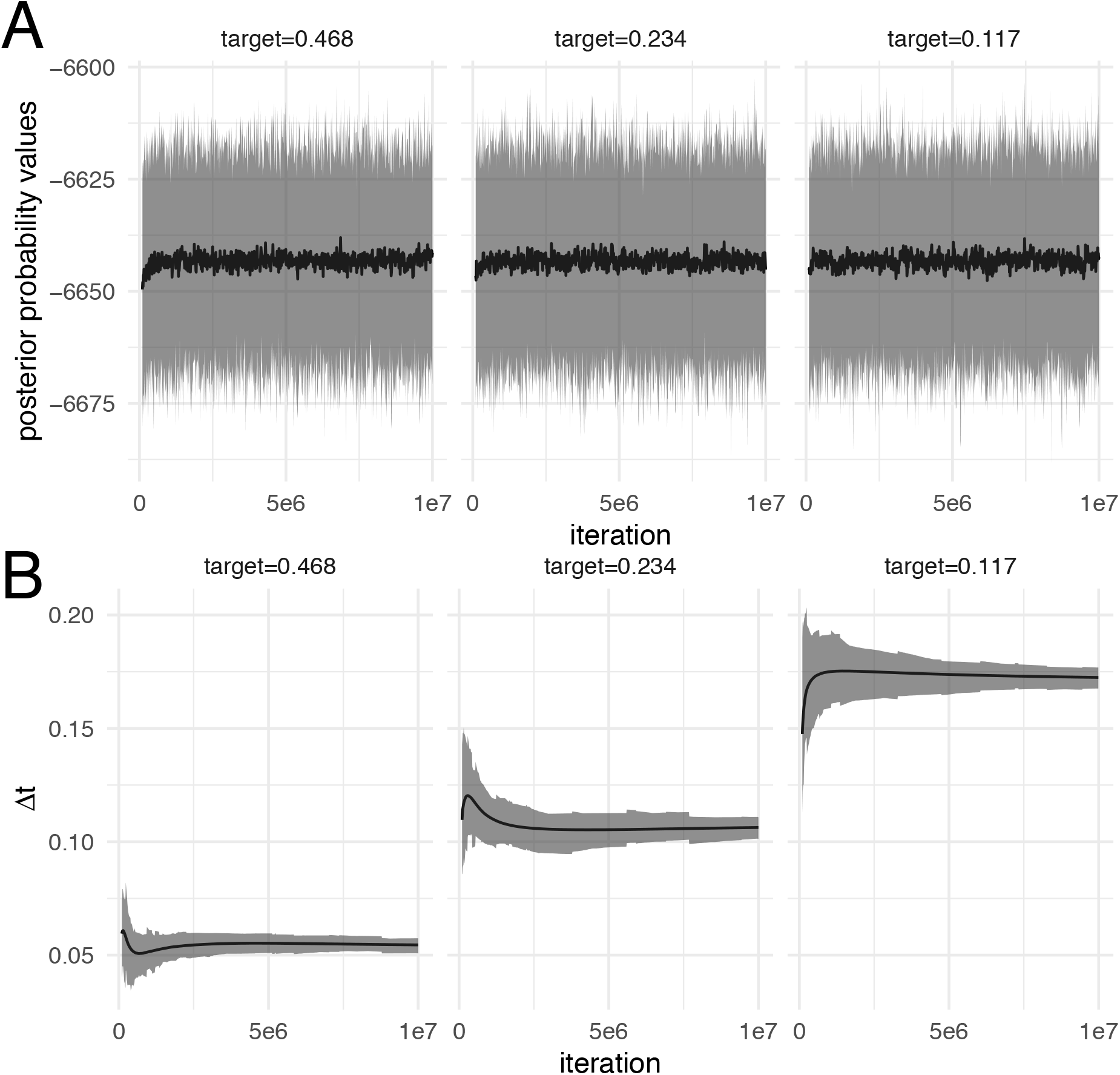
Distribution of posterior probability values at different iterations over 100 analyses. **A** The black line denotes the mean posterior probability estimates (y-axis) over 100 analysis at different iterations (x-axis). The grey area denotes the 95% highest posterior density interval of posterior probability estimates over these 100 analyses at different iterations. The different subplots show the results using runs with 3 different target acceptance probabilities, leading to different temperature differences between the chains. **B** The black line denotes the mean temperature difference Δ*t* between chains on the y-axis over 100 analyses at different iterations on the x-axis. The grey area denotes the 95% highest posterior density interval of Δ*t* over these 100 analyses at different iterations.

**Figure 2:**
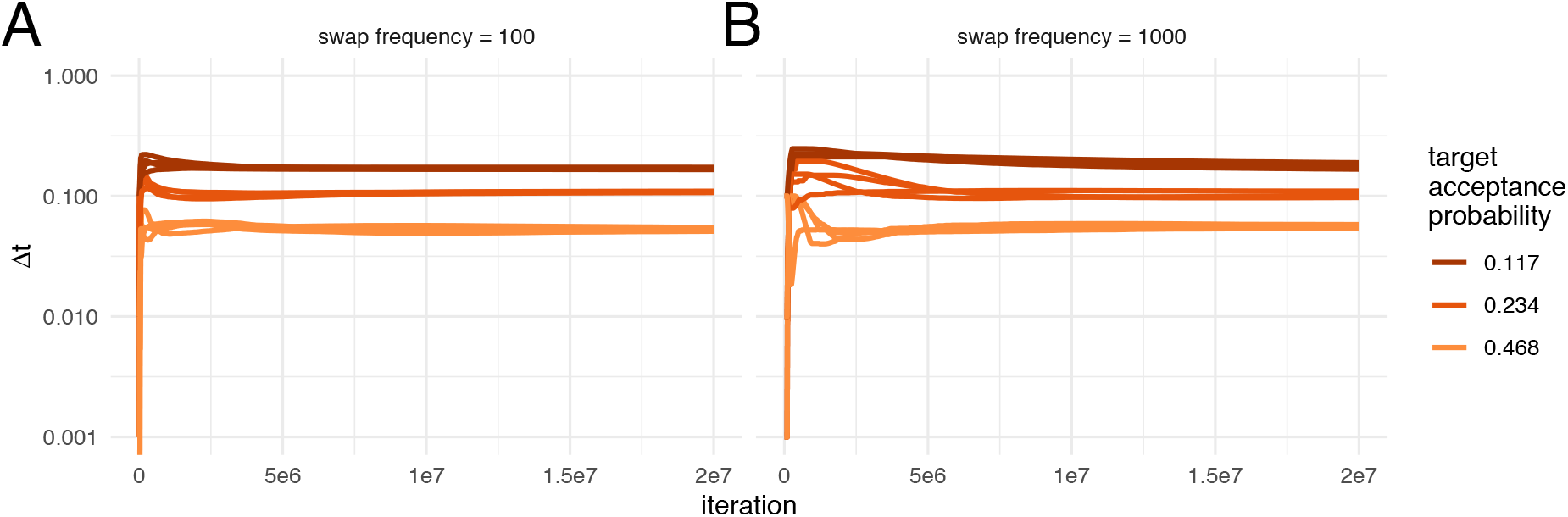
Automatic tuning of the temperature to achieve different acceptance probabilities. Here we show how the temperature difference between chains (y-axis) is adapted during the course of an adaptive parallel tempering run on the x-axis. Each color represents runs with different target acceptance probabilities. For each of the four different target acceptance probabilities, we started runs at four different initial temperatures. **A** Acceptance probability over the course of a run when swaps of states between chains are proposed every 100 iteration. **B** Acceptance probability when swaps are proposed every 1000 iteration.

### The effect of heating on exploring the posterior

In order to explore how heating affects exploring the posterior probability space, we next compared effective sample size (ESS) between regular and parallel tempering at different temperatures on a dataset where we do not expect any problems in exploring the posterior space caused by several local optimas. ESS values denote the number of effective samples if all samples would be drawn randomly from a distribution and are estimate here using Tracer (Rambaut et al., 2018).

To compare ESS values, we ran the Bayesian coalescent skyline (Drummond et al., 2005) analysis of Hepatitis C in Egypt (Ray et al., 2000) for 4 * 10^7^ iterations using MCMC in 100 replicates. Additionally, we performed 100 replicates using parallel tempering with 4 different chains for 1 * 10^7^ iterations using 3 different target acceptance probabilities, 0.468, 0.234 and 0.117. The different chain lengths between MCMC and parallel tempering are chosen such that the overall number of iterations over the cold and heated chains is the same for parallel tempering as for MCMC. After running all 4 times 100 analyses, we computed the ESS values of the posterior probability estimates using loganalyser in BEAST 2 (Bouckaert et al., 2014).

As shown in figure 3A, the average ESS values are highest for the cold scenario when using parallel tempering and decrease with lower target acceptance probabilities. Lower target acceptance probabilities mean lower higher temperatures of heated chains in those analyses.

**Figure 3:**
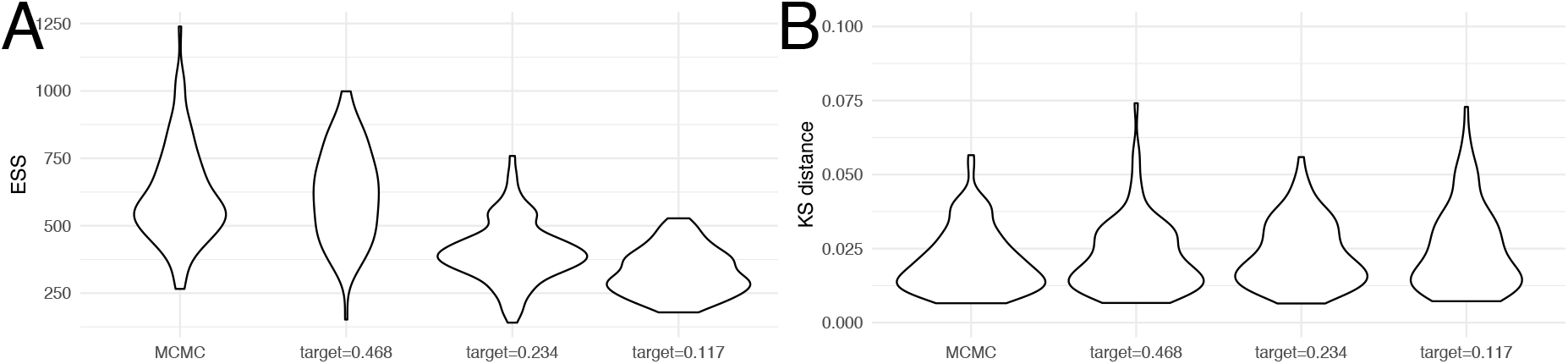
Convergence of coupled MCMC and regular MCMC using posterior ESS values and Kolmogorov Smirnov distances. **A** Here, we show the distribution of posterior ESS values after 4 * 10^7^ for regular MCMC and after 1 * 10^7^ for parallel tempering with 4 chains, so wall time for MCMC runs was much larger than for parallel tempering. When running the analyses with parallel tempering, we used 3 different target acceptance probabilities. **B** Here we show the distribution of Kolmogorov Smirnov distances between individual runs and the concatenation of all individual runs. We assume that all 400 runs concatenated describe the true distribution of posterior values and then take the KS distance as a measure of how good an individual run approximates that distribution. The smaller a KS value, the better the true distribution is approximated.

We next tested if higher ESS values actually correspond to a run approximating the distribution of posterior probability values better. To do so, we compared Kolmogorov-Smirnov (KS) distances between individual runs and the true distribution of posterior values. The KS distance denotes the maximal distance between two cumulative density distributions, which is smaller the better to distributions match. Since we can not directly calculate the true distribution of posterior values, we concatenated the 400 regular and parallel tempering runs and used the concatenated distribution of posterior values as the true distribution. Figure 3B shows the distribution of KS distances between individual runs using regular and parallel tempering to what we assume to be the true distribution. In contrast to the comparison of ESS values, we find that the distribution of KS distances is fairly comparable across all methods. This indicates that in this analysis, parallel tempering with 4 individual chains performs equally well as MCMC run for 4 times as long. It also shows that the differences in ESS values between the parallel tempering runs with different target acceptance probabilities are indicative of more swaps, rather than a better approximation of the true posterior probability distribution.

We next compared the inference of trees on a dataset DS1 that has proved problematic for tree inference using MCMC (Lakner et al., 2008; Höhna and Drummond, 2011; Maturana Russel et al., 2018). This dataset has many different tree island, transitioning between which is highly unlikely due to very unfavourable intermediate states (Hohna and Drummond, 2011).

**Figure 4:**
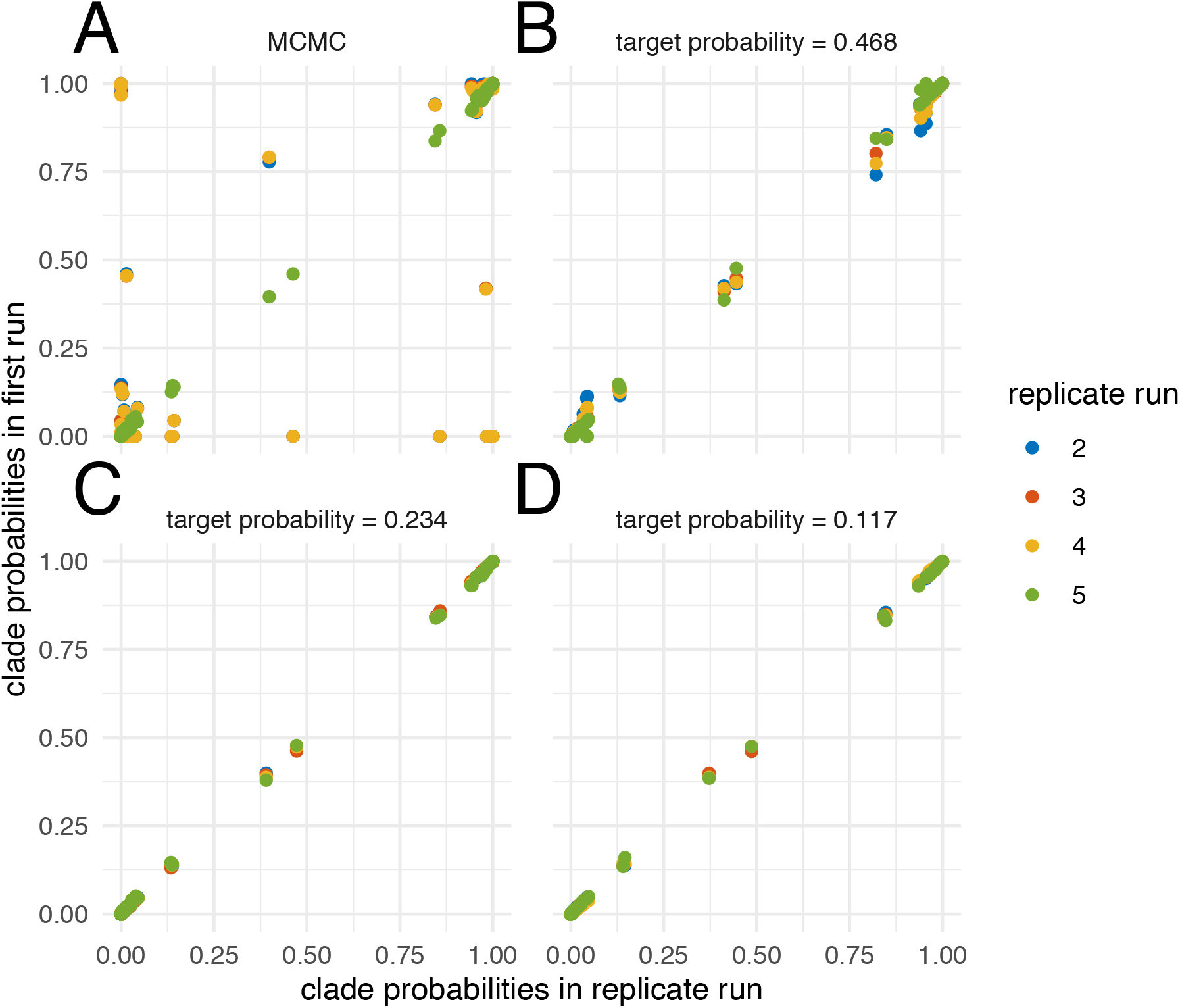
Inferred clade probabilities between different replicate runs. Here we compare inferred clade credibilities between one run (y-axis) and four replicates from different starting points (x-axis) using MCMC **A** and adaptive parallel tempering run with target acceptance probabilities of 0.468 **B**, 0.234 **C** and 0.117 **D**.

We ran the dataset using MCMC for 5 * 10^7^ iteration and parallel tempering for 5 * 10^7^ with 4 different chains. We ran parallel tempering targeting three different acceptance probabilities, that is 0.117, 0.234 and 0.468. MCMC gets stuck in different local optimas, resulting in differences between inferred clade credibilities across different runs (see figure 4). The clade credibilities are more comparable when targeting an acceptance probability of 0.468 and become more consistent between the different runs with acceptance probabilities of 0.234 and 0.117. At higher target acceptance probabilities (i.e. lower temperatures), the heating of chains is not sufficient to efficiently travel between local optimas.

We additionally compared how well the different runs approximate the posterior probability distribution compared to how long they ran. Several MCMC runs sample from a different posterior probability distribution compared to parallel tempering with a low target acceptance probability and a high temperature (see figure S5). When running parallel tempering with a relatively high target acceptance probability of 0.468, the KS distance to the reference distribution decreases relatively slowly with the number of iterations compared to lower acceptance probabilities. This suggests that at lower temperatures (i.e. higher acceptance probabilities), some of the chains get stuck in local optimas.

## Conclusion

Next generation sequencing has lead to ever larger datasets of genetic sequence being available to researcher. To study these, more and more complex models are developed, many of which are implemented in the Bayesian phylogenetic software platform BEAST2 (Bouckaert et al., 2014). Parallelising these models can often be hard or even impossible and MCMC analyses often have to be run on single CPU cores.

Alternatively, parallel tempering can make use of multiple cores, but a full featured version was so far not available in BEAST 2. Parallel tempering, however, requires choosing optimal temperatures of heated chains. We here circumvent the issue of choosing optimal temperatures by adaptively tuning the temperature difference between heated chains to achieve a target acceptance probability implemented for BEAST 2.5 (Bouckaert et al., 2019). In order to only have one parameter to tune, we assume that the temperature difference between heated chains is geometrically distributed and only tune the temperature difference between those. We show that this adaptive tuning of the temperature difference is targeting different acceptance probabilities well, starting from various different initial values. Alternatively, the temperature differences could be defined between individual chains, which would require tuning the number of chains minus 1 temperatures (Miasojedow et al., 2013). While potentially leading to a more optimal spacing of temperatures between individual heated chains, we here chose an approach where the number of parameters that have to be tuned is minimal. We hope that this minimizes the amount of tuning needed and reduces the complexity of setting up an analysis to the same level as for a regular MCMC analysis and therefore makes it as user friendly as possible.

We next compared convergence between using different target acceptance probabilities as well as regular MCMC. We find that ESS values are comparable between parallel tempering with *N* chains and a relatively high target acceptance probability of 0.468 and regular MCMC that ran N times longer. ESS values decreased on this dataset when using lower target acceptance probabilities and therefore higher temperatures. When comparing how well the true posterior distributions are approximated between the different target acceptance probabilities, we found that using different target values did not significantly influence how well the distributions are approximated.

ESS values are estimated by computing the auto-correlation time between samples. We suspect that swapping the states between chains strongly decreases this autocorrelation. In turn, this would mean that the more frequently states are exchanged, the shorter this auto-correlation become, which would increase ESS values.

This indicates that convergence statistics like the scale reduction factor (Brooks and Gelman, 1998), might be better suited to assess convergence than ESS values. Since the parallel tempering runs required *N* times fewer iterations of the cold chain to approximate the distribution of posteriors values as well, parallel tempering can help speed up analysis by a factor *N* that can be chosen to be proportional to the number of CPU’s used.

The adaptive parallel tempering algorithm is compatible with other BEAST 2 packages and therefore works with any implemented model that does not directly affect the MCMC machinery. This will help analyzing larger datasets with more complex evolutionary and phylodynamic models without requiring additional user specifications other then the number of heated chains.

## Acknowledgement

NFM is funded by the Swiss National Science foundation (SNF; grant number CR32I3_166258). RRB is supported by by Marsden grant 18-UOA-096 from the Royal Society of New Zealand.

## Authors contribution

NFM and RB implemented the code, NFM performed the analyses and NFM and RB wrote the paper.

