## Supplement for "Adaptive parallel tempering for BEAST 2"

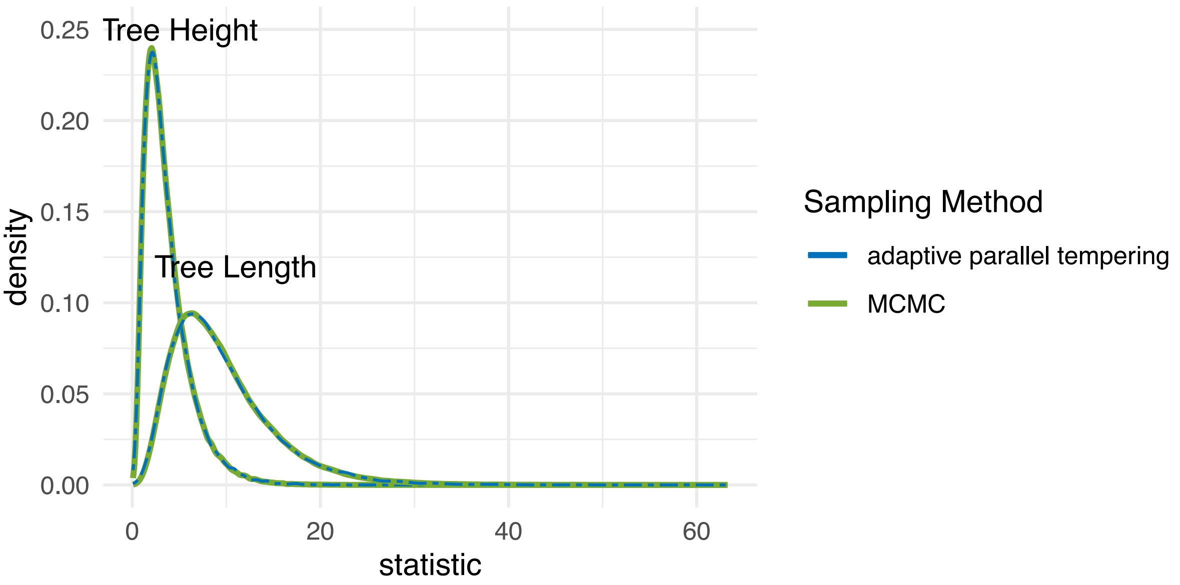


Figure S1: **Comparison of inference between adaptive parallel tempering and regular MCMC**. Comparison of the distribution of tree heights and tree lengths sampled under the structured coalescent using MultiTypeTree. The inferred distribution of tree heights and tree lengths match up between MCMC and the cold chain in parallel tempering.

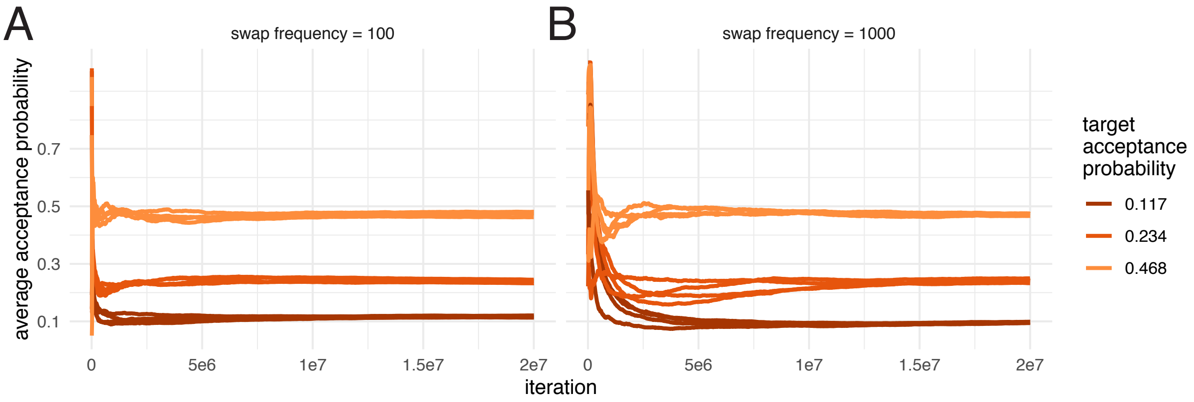


Figure S2: **Acceptance probabilities over the course of an analyses with different target acceptance probabilities.** Here we show the global acceptance probability during the course of an adaptive parallel tempering run on the x-axis. Each color represents runs with different target acceptance probabilities. For each of the four different target acceptance probabilities, we started runs at four different initial temperatures. **A** Acceptance probability over the course of a run when swaps of states between chains are proposed every 100 iteration. **B** Acceptance probability when swaps are proposed every 1000 iteration.

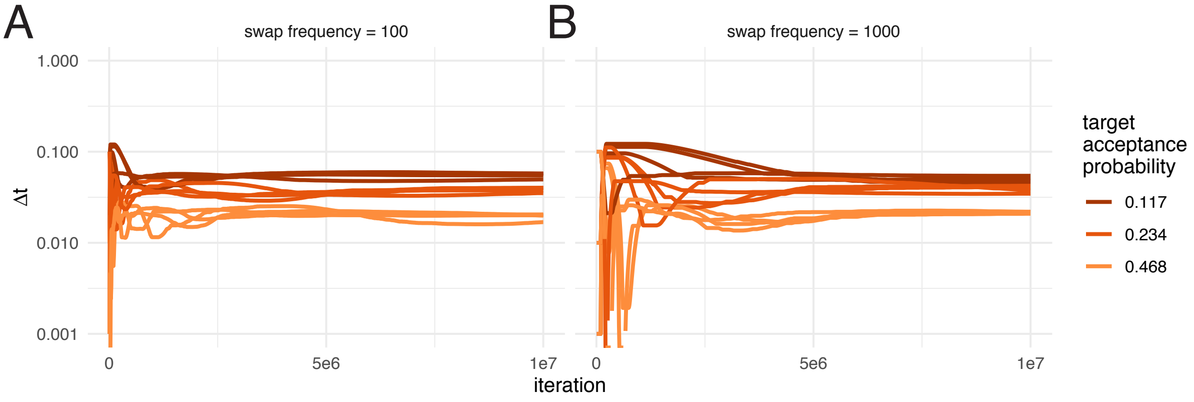


Figure S3: **Automatic tuning of the temperature to achieve different acceptance probabilities for a MASCOT analysis of influenza A/H3N2.** Here we show how the temperature difference between chains (y-axis) is adapted during the course of an adaptive parallel tempering run on the x-axis. Each color represents runs with different target acceptance probabilities. For each of the four different target acceptance probabilities, we started runs at four different initial temperatures. **A** Acceptance probability over the course of a run when swaps of states between chains are proposed every 100 iteration. **B** Acceptance probability when swaps are proposed every 1000 iteration.

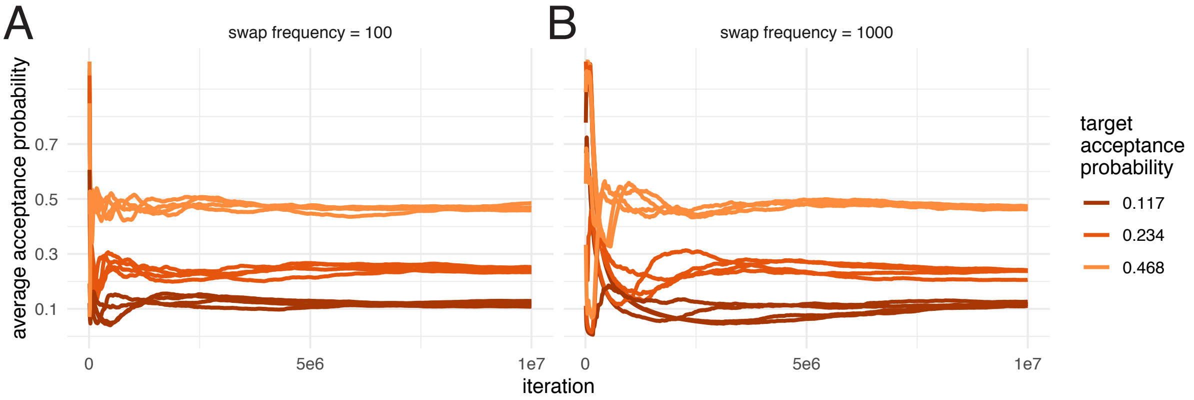


Figure S4: **Acceptance probabilities over the course of an analyses with different target acceptance probabilities for a MASCOT analysis of influenza A/H3N2.** Here we show the global acceptance probability during the course of an adaptive parallel tempering run on the x-axis. Each color represents runs with different target acceptance probabilities. For each of the four different target acceptance probabilities, we started runs at four different initial temperatures. **A** Acceptance probability over the course of a run when swaps of states between chains are proposed every 100 iteration. **B** Acceptance probability when swaps are proposed every 1000 iteration.

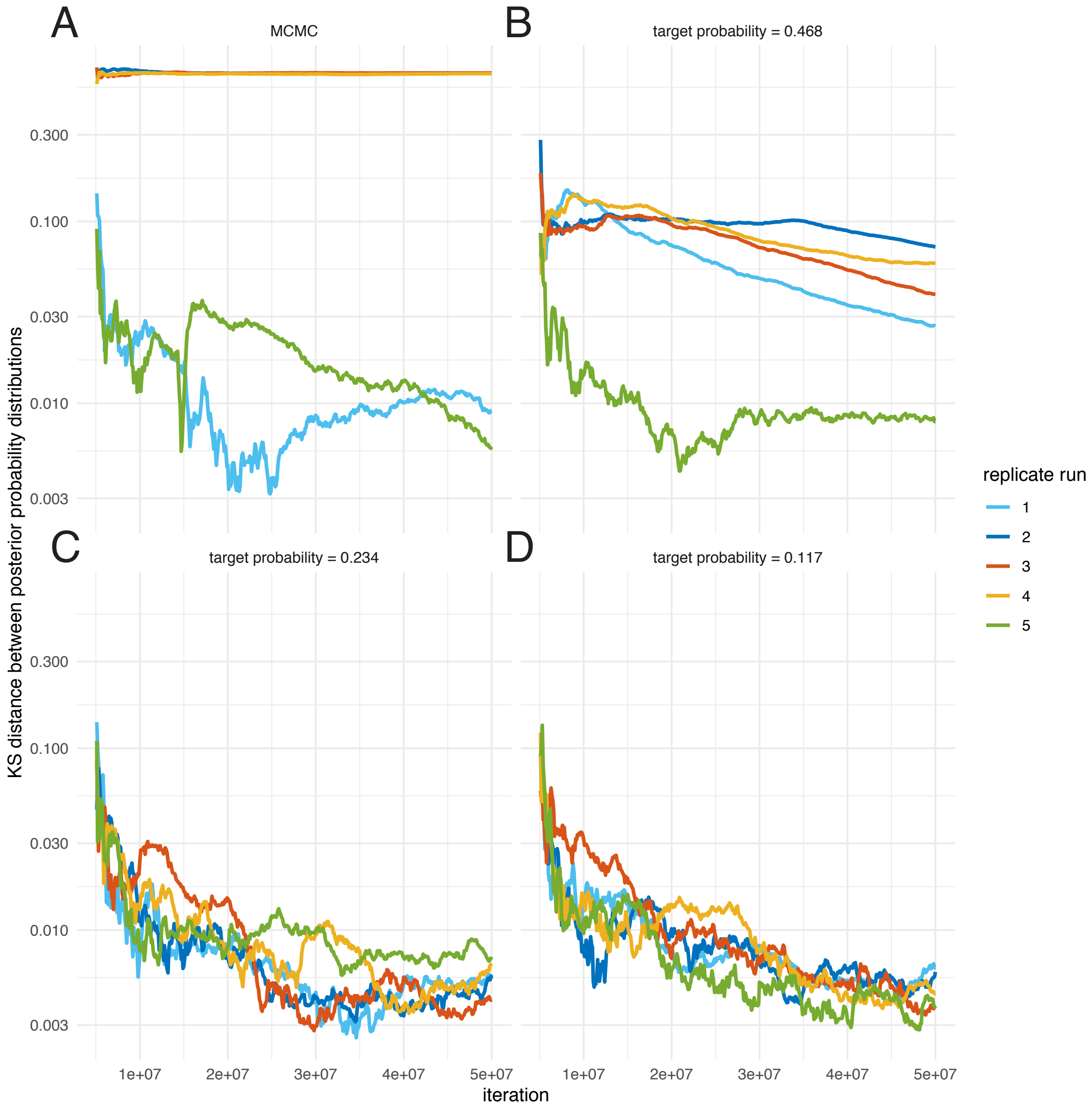


Figure S5: **Convergence versus number of iterations of the posterior probability distribution for different acceptance probabilities.** Here, we show the Kolmogorov-Smirnov (KS) distance between the inferred posterior probability distribution and a reference posterior probability distribution on the y-axis up to the iteration on the x-axis. The different plots show the KS distance over the number of iterations for MCMC (**A**), and parallel tempering with an target acceptance probability of 0.468 **B**, 0.234 **C** and 0.117 **D**. The reference distribution consists out of 4 of the 5 runs with a target acceptance probability of 0.117. For each run with any target acceptance probability and with MCMC, the corresponding run is removed from the reference distribution.
